# Parallels and divergences in landscape genetic and metacommunity patterns in zooplankton inhabiting soda pans

**DOI:** 10.1101/2022.12.17.520856

**Authors:** Zsófia Horváth, Tsegazeabe Hadush Haileselasie, Csaba F. Vad, Robert Ptacnik, Luc De Meester

## Abstract

Ecological processes maintaining landscape genetic variation and metacommunity structure in natural landscapes have traditionally been studied in isolation. Their integrated study may hold important information as to what extent the effect of major ecological processes are species-or landscape-specific, resulting in a more coherent picture on the spatial organization of biodiversity. Here, we explicitly compared the relative importance of spatial and environmental drivers of both cladoceran metacommunity structure as well as landscape genetic structure of its most widespread member, the water flea *Daphnia magna*, in soda pans of the Seewinkel region in Austria. This landscape of soda pans is characterized by strong environmental gradients and unidirectional wind acting as a key dispersal agent among these temporary habitats. Our study shows both parallels and divergences in the relative importance of local environmental sorting and spatial connectivity in determining landscape genetic versus metacommunity structure. The metacommunity is structured primarily by the environment, while in the *D. magna* metapopulation, the spatial signal is predominant. The much weaker environmental signal in *Daphnia* can be explained by the fact that the microsatellite markers are presumably neutral and was confirmed by a per-allele analysis. An important parallel between metacommunity and landscape genetic structure is the strong signal of the prevailing wind direction in determining the spatial pattern. This suggests that for both community assembly in cladocerans and population assembly in *D. magna*, wind plays an important role in determining connectivity among soda pans, thereby affecting dispersal and colonization rates, influencing both local species and genetic composition.

## Introduction

Dispersal is essential for maintaining connectivity among local populations and communities. It drives landscape-level heterogeneity in genetic and species composition, in interaction with other processes including local environmental sorting (selection) and drift (Vellend and Geber 2005). This implies that at least theoretically, metapopulations and metacommunities are in essence structured by the same processes (Vellend 2016). Drivers of landscape genetic structure are indeed expected to show strong parallels to the drivers of variation in species composition in local communities across landscapes (cf. metacommunity structure; Leibold et al. 2004). In both cases, dispersal is linked to landscape connectivity, and can lead to a pattern of increasing similarities with decreasing geographic distance (i.e., distance decay of similarity). Similarly, in both cases, environmental heterogeneity can structure patterns of convergence and divergence. Vellend (2010, 2016) highlighted the parallels between metacommunity (species sorting, dispersal, neutral dynamics and speciation) and metapopulation processes (natural selection, gene flow, neutral dynamics and mutation). Yet, due to a lack of explicit comparative studies in the same landscapes, little is known on the relative contribution of these processes to shaping landscape genetic variation and species composition.

The explicit comparative test of the drivers of landscape genetic and metacommunity structure requires studies performed in the same landscape with the same statistical tools. In metacommunity ecology, several analyses have been developed to disentangle spatial from environmental effects (Borcard et al. 1992; Peres-Neto et al. 2006), followed by numerous empirical studies that tested the relative role of environmental sorting and dispersal on biodiversity and community composition. Landscape genetics similarly involves testing for the relative importance of local environment and dispersal on genetic metapopulation structure and gene flow (Manel *et al*. 2003; Guillot *et al*. 2005). While population genetic studies traditionally focused on geographic distance as a key structuring factor (isolation-by-distance; Bohonak 1999, 2002), there is increasing evidence for strong environmental gradients mediating effective gene flow and structuring landscape genetic variation (Nosil et al. 2005; Orsini et al. 2013; Sexton et al. 2014; Moncada et al. 2021). Given these parallels, the same multivariate approaches that are applied in metacommunity ecology can deliver information on the relative importance of spatial and environmental predictors for explaining not only community composition in a landscape, but also landscape genetic structure (Orsini et al. 2013). While there is an increasing interest in multi-species approaches (Manel and Holderegger 2013; Wang and Bradburd 2014; Hein et al. 2021) resulting in a growing number of multi-species landscape genetic studies (Manier and Arnold 2006; Goldberg and Waits 2010; Frantz et al. 2012; Richardson 2012), explicit comparisons of metacommunity and metapopulation level patterns are largely lacking.

Here, we compare metacommunity structure of cladoceran zooplankton to landscape genetic variation in its dominant member, the water flea *Daphnia magna*, in the same set of habitats, saline temporary waters (soda pans) in the Seewinkel region of Austria (**Fig. 1**). Specifically, we aim to explicitly test the relative strength of major structuring forces (i.e., environment and space) across the two levels of ecological organization. Our study landscape features strong environmental gradients of salinity and turbidity that are expected to act as strong selective forces for the local biota (Horváth et al. 2014). At the same time, the flat lowland area experiences strong unidirectional winds, which is expected to result in a directional spatial imprint on passively dispersing organisms (Epele et al. 2021; Kling and Ackerly 2021). The latter was documented for the metacommunity structure of zooplankton in an earlier study (Horváth et al. 2016). Soda pans therefore represent an attractive model for a simultaneous analysis of landscape genetic and community variation along strong environmental and connectivity gradients within a relatively small region.

**Figure 1.**
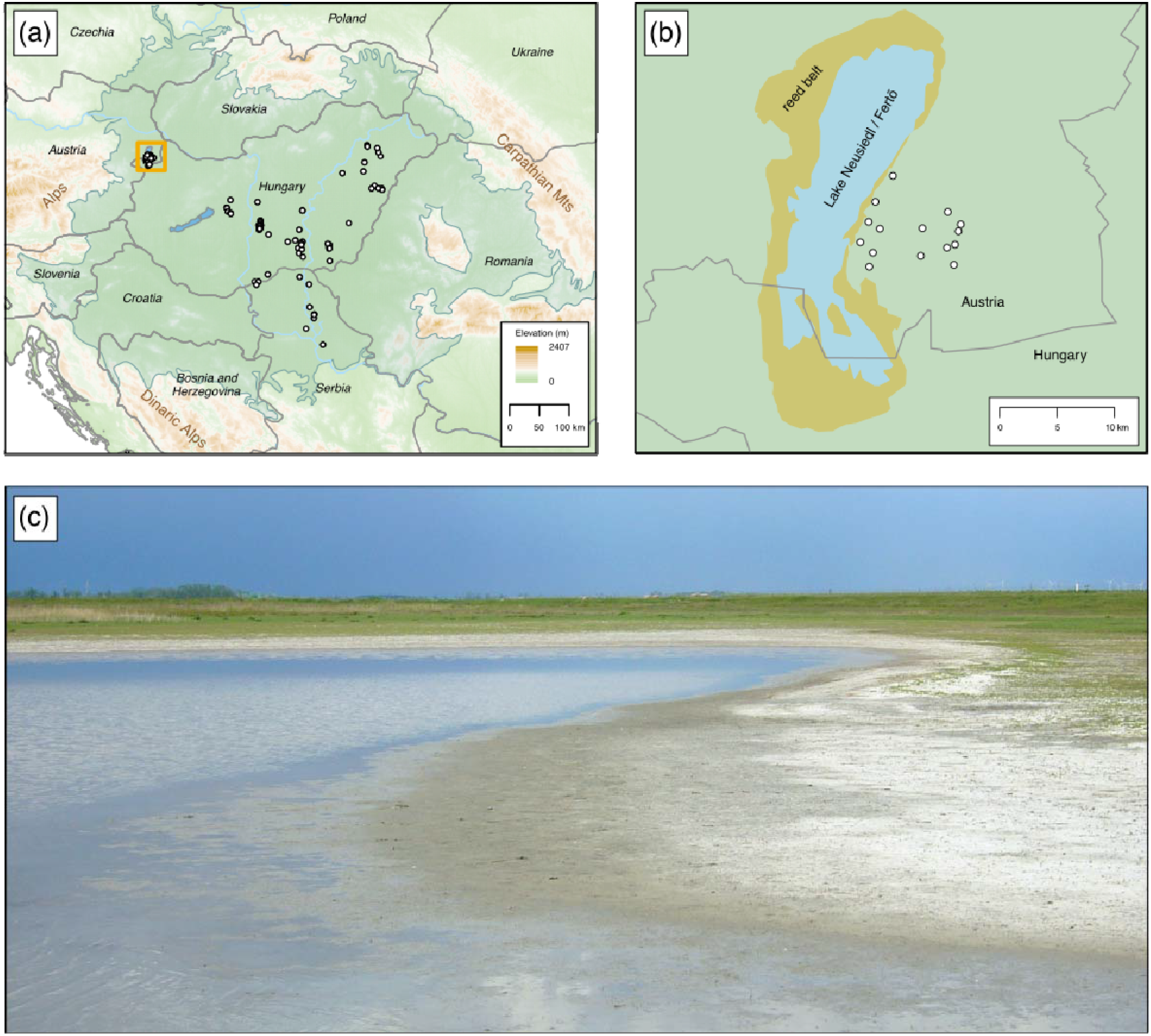
– (a) Map showing the Pannonian Plain as an enclosed lowland between the Alps and the Carpathian Mountains in East-Central Europe, where dark green shading indicates the Pannonian biogeographic region (white circles: soda pans). The orange frame on the border between Hungary and Austria highlights the study region (Seewinkel). (b) The studied 15 soda pans in the Seewinkel region (white circles). (c) An example of the study habitats during the survey.

## Materials and Methods

### Study area and sample collection

Soda pans are shallow, saline and temporary fishless waterbodies, which generally dry out in summer. Within Europe, their occurrence is restricted to the Pannonian Plain; the lowlands of Hungary, eastern Austria, and northern Serbia (**Fig. 1a**; (Boros et al. 2014). Soda pans usually have a vegetation-free lakebed and no or only scarce emergent vegetation along the shorelines (**Fig. 1c**). The pans are alkaline and their salinity exhibit a wide gradient from almost freshwater to hypersaline conditions (Boros et al. 2014). They are generally well-mixed and not stratified due to their exposure to wind and their high surface to depth ratio. Due to the absence of fish and the hypertrophic conditions, zooplankton communities are bottom-up regulated (Horváth et al. 2013, 2014). The predominant environmental gradient, salinity, is a strong selection factor, and only a limited set of highly tolerant species account for the largest part of zooplankton biomass (Horváth et al. 2014). This includes *Daphnia magna*, a widely-used model species in landscape genetic studies, which is at the same time the most widespread species in these habitats (Tóth et al. 2014).

According to a recent survey (Tóth et al. 2014), there are 23 well-preserved soda pans in the Seewinkel region of Austria. We collected zooplankton samples from all 23 pans in spring 2014 (28-29 April), by randomly collecting 20 l of water from the central part of each habitat. Water was then sieved through a plankton net with a mesh size of 45 μm and preserved in 70% ethanol on the field. In the lab, we applied subsampling to count the abundance of Cladocera species in the samples; counting and identifying at least 300 individuals represents a subsample of 0.5-15%, depending on the total density in the sample. Afterwards, the entire sample was checked for rare species. We also isolated either all or at least 20 *D. magna* individuals, and preserved them in 96% ethanol until genotyping.

We determined the GPS location of the approximate centre of the pans, and measured water depth (Z), conductivity, and pH in the field. Water samples were subsequently analysed in the lab to quantify the concentrations of total suspended solids (TSS), total phosphorus (TP) and chlorophyll *a* (Chl). TSS was measured gravimetrically by filtering water (1–25 ml, depending on turbidity) through pre-dried (oven-drying at 105 °C) and pre-weighted glass fibre filters (GF/F). TP was measured by using persulfate digestion (Clesceri et al. 1999), followed by the ascorbic acid colorimetric method (Hansen et al. 1999). For Chl, we used fluorometry with acetone extraction (Arar and Collins 1997), without correcting for phaeophytin.

During our sampling campaign, 15 out of the 23 pans were found to host *D. magna* populations (**Fig. 1b**, **Table S1** in Supporting information). We therefore decided to use only these 15 sites in our comparative landscape genetic and metacommunity analysis. In our landscape genetic analysis, we thus capture the complete metapopulation of this species inhabiting soda pans.

### Population genetic analysis: DNA extraction and microsatellite amplification

We screened for variation at 13 polymorphic neutral markers (see **Table S1** in Supporting information; based on a set of 84 primers used by Orsini et al. 2012) in all 15 soda pan *D. magna* populations. On average, 19 animals per population were genotyped (**Table S1**). We genotyped 20 individuals from 11 pans; in the remaining four pans 21, 19, 16 and 10 individuals were genotyped. As no more animals were available in our samples, these latter numbers could not be increased any further.

Genomic DNA was extracted from individual *Daphnia* using the HotShot method (Truett et al. 2000; Montero-Pau et al. 2008). DNA amplifications were carried out in a final reaction volume of 11.5 µl containing 1.5 µl of template genomic DNA and 10 µl of a mixture of 0.2-0.3 µM of each primer (labelled with fluorescent dyes, 8 for multiplex M01 and 5 for M03) and 1x Qiagen Multiplex PCR Master Mix buffer (Qiagen®). PCR amplification cycling conditions included an initial denaturation step at 95 °C for 15 minutes, 30 cycles of denaturation at 94 °C for 30 seconds, annealing at 54 (for multiplex M03) or 56°C (for multiplex M01) for 1.5 minutes, extension at 72 °C for 1.5 minutes, and a final elongation step at 60 °C for 30 minutes.

Amplified PCR products were separated on an ABI3130 capillary sequencer (Applied Biosystems®). Allele sizes (fragment lengths) were determined by comparison with an internal Liz500 size standard using the software GeneMapper (v3.7; Applied Biosystems®).

## Data analysis

### Population genetic structure

Standard measures of genetic variation, including percent of polymorphic loci, total (A) and mean number of alleles per locus (allelic richness, A_r_), expected heterozygosity (H_e_), observed heterozygosity (H_o_), inbreeding coefficients (F_IS_: individuals relative to the subpopulation; F_IT_: individuals relative to the total population) and fixation index (F_ST_, quantifying subpopulation differentiation compared to the total population) were calculated in R (R Development Core Team 2012) using packages “adegenet” (Jombart 2008), “polysat” (Clark and Jasieniuk 2011), and “pegas” (Paradis 2010). A locus was considered polymorphic if the most common allele had a frequency less than 0.99. Deviations from Hardy-Weinberg equilibrium were tested with Monte Carlo permutation tests (Goudet’s G-statistic with 1000 permutations) in “pegas” (Paradis 2010); these tests were carried out separately for all loci and populations.

The extent of genetic differentiation among populations was assessed by comparing pairwise F_ST_ values (Weir and Cockerham 1984) across all loci, using the package “hierfstat” in R (Goudet 2005). A global F_ST_ value for the region across all loci and populations was calculated with the package “polysat” (Clark and Jasieniuk 2011), and its significant difference from zero was assessed using 1000 permutations.

### Genetic and community matrix

For obtaining a data matrix representing genetic differentiation, we first built a PCoA on the pairwise F_ST_ values (based on Bruvo distances) with the package “ape” for R (Paradis et al. 2004). In cyclically parthenogenetic organisms such as the water flea *D. magna*, clonal structure might complicate genetic analyses, as multiple individuals might belong to the same clone. In the soda pan populations, however, all sequenced individuals were characterized by different multilocus genotypes, and this allowed us to use data on allelic frequencies in the PCoA analysis. In a second step, we used the scores of all PCoA axes as the dependent matrix in the subsequent variation partitioning analyses.

For the metacommunity data, we also ran a PCoA, this time based on Bray-Curtis dissimilarities of the double square root transformed abundance data of the Cladocera communities (after the exclusion of two singletons), and then extracted the scores of all PCoA axes.

### Environmental matrix

Among the environmental predictors, we log-transformed the non-normally distributed ones (Z, TSS, Chl, TP). In soda pans, strong correlations between environmental variables are frequent (Horváth et al. 2014, 2016); hence we first inspected pairwise Pearson’s correlations among our predictors (**Table S2** in the Supporting information). Based on that, we performed an *a priori* data reduction by excluding TP from our initial set of explanatory variables (because it was strongly correlated to almost all other predictors), and only retained Z, TSS, Chl, and conductivity. These four predictors were then standardised to zero mean and unit variance.

To select the most relevant environmental predictors, we applied an automatic stepwise model selection from an initial RDA (Redundancy Analysis) model including the four standardised environmental explanatory variables and the PCoA axes scores (either resulting from the metapopulation or the metacommunity) as the dependent matrix. We selected for significant predictors by using a permutation test (“ordistep” function in “vegan”, with direction=“both” and 2000 permutations; Oksanen et al. 2018).

### Spatial matrix

For quantifying spatial predictors, we employed spatial eigenvector methods in order to get a more explicit representation of spatial patterns. Because of the strong prevalent wind direction (Horváth et al. 2016), we applied both Moran’s Eigenvector Maps (MEM; Dray et al. 2006), as well as Asymmetric Eigenvector Maps (AEM; Blanchet et al. 2008), to identify patterns both with (AEM) and without taking directionality into account (MEM). We used the same connectivity matrix (binary neighbours list) for both sets of eigenvectors, built with a threshold larger than the longest distance between any pair of pans (to enable all possible connections between pans). For the AEM eigenvectors, the dominant wind direction was added as the vector angle (292.5°) of the directional component (see also **Fig. S1** in Supporting information). MEM and AEM eigenvectors were computed with the “spdep” (Bivand et al. 2015) and “adespatial” packages (Dray et al. 2018). To select the most relevant eigenvectors, we applied automatic stepwise model selection in a similar manner as for environmental predictors.

### Relative role of environmental and spatial predictors

To quantify the relative importance of environmental and spatial predictors, we carried out variation partitioning analyses with the “vegan” package (Oksanen et al. 2018). Here, the PCoA axes scores (either resulting from levels of genetic differentiation or community dissimilarity) were used as the dependent matrix, together with a selected set of environmental and spatial predictors.

In order to make results the most comparable across the metapopulation and the metacommunity, we used a merged set of predictors, by including all predictors that proved significant for at least one of the dependent matrices. For the metacommunity, the only significant environmental predictor was TSS, while for the metapopulation, no significant or marginally significant predictor was found. Hence we used TSS as the environmental predictor for both datasets (in all four analyses). In the case of AEM eigenvectors, one eigenvector (AEM 10) was significant for the metacommunity, and another (AEM 3) was marginally significant for the metapopulation. Therefore, we included both in our variation partitioning models, to use the exact same set of variables for both datasets. By doing so, we can ensure that lower variation explained by either environment or space in our comparative analysis does not arise from a difference in the number of predictors considered.

### Partitioning analyses on single species and alleles

To explore variation in the spatial and environmental signal in the individual community members and verify whether the degree of dispersal limitation as observed for individual species might be linked to their occupancy in the metacommunity, we ran a series of variation partitioning analyses on each community member (excluding singletons).

For the analyses at the level of individual species, we used the double square root transformed community matrix (abundance data) after the exclusion of the two singletons. For individual species, we applied automatic stepwise model selection on linear models. We selected for significant predictors by using a stepwise model selection by AIC (“stepAIC” function in “MASS”, with direction=“both” and 2000 permutations; Ripley et al. 2013).

Given that for the five species, different environmental predictors and eigenvectors proved important (see **Table S3** and **S4** in Supporting information), we decided not to unify predictors as carried out above for genetic and community data, and used the respective sets of significant predictors for each species. The resulting values of pure environmental and spatial effects (**Table S5** in Supporting information) were related to the regional occupancy of species using linear models to test the idea that dispersal limitation is more apparent and species sorting less effective in rare than in common species (Horváth et al. 2019). To ensure that patterns are not due to any sampling effect (Leibold and Chase 2017; Viana and Chase 2019), we also tested the relationship between total explained variation and regional occupancies.

To explore whether a similar pattern exists for landscape genetic structure of the dominant community member, we also ran such analyses for genetic variation within the *D. magna* metapopulation. We could not run such an analysis at the level of single genotypes because all sequenced individuals in our study were characterized by different multilocus genotypes. We thus decided to carry out these analyses at the level of alleles, checking whether spatial signals are more important in less common compared to more widespread alleles, quantified as the number of occurrences in the 15-site landscape. This linear regression involved all alleles after the exclusion of singletons. We did not unify predictors, because the high number of variation partitioning analyses (one for each of the 131 non-singleton alleles) resulted in the full set of predictors being selected at least once. We report the number of times a given environmental or spatial predictor was selected in the 131 models in the Supporting information (**Fig. S2**).

### Isolation-by-distance

To test whether genetic differentiation among populations increases with geographic distance, we carried out a Mantel test on pairwise measures of genetic differentiation (F_ST_/(1 − F_ST_)), against pairwise geographic distances with 2000 permutations. Given the expectation that dispersal rates might be higher in the direction aligned with the wind, we carried out two Mantel tests, decomposing geographic distances into a component parallel to the predominant wind direction and a component orthogonal to the predominant wind direction (see also **Fig. 4b**).

## Results

The cladoceran metacommunity counted a total of seven species. Apart from two singletons (*Alona rectangula* and *Megafenestra aurita*), the other species all occurred in at least three habitats, with *D. magna* being the most frequent species among them (occurring in all 15 habitats). The other species were *Moina brachiata* (13), *Macrothrix hirsuticornis* (8), *Chydorus sphaericus* (5), and *Daphnia atkinsoni* (3). Mean species richness (± SD) in the habitats was 3.07 ± 1.22.

Across the 13 loci, a total of 158 alleles were detected (with a mean ± SD of 12.2 ± 4.2 alleles per locus and 5.7 ± 2.6 alleles per locus in a population) in *D. magna* from the soda pans (**Table S1** in Supporting information). 27 alleles were singletons while 131 alleles occurred in at least two habitats. The regional mean allelic richness was 6.08 ± 3.2 (range: 2-22 alleles per locus in a population, **Table S1** in Supporting information). All loci were polymorphic. Summaries of genetic diversity parameters are presented in **Table S1** (for each habitat) and **Table S6** (for each locus) in the Supporting information, together with the pairwise F_ST_ values of all habitat pairs (**Table S7**). According to Monte Carlo tests, the observed level of heterozygosity was significantly lower than what we expect under Hardy-Weinberg equilibrium both for the populations (**Table S1** in Supporting information) and for the individual alleles (**Table S6** in Supporting information). Global genetic differentiation among populations was significant (F_ST_=0.12, t=211.56, df=999, p<0.001).

While symmetric MEMs did not reveal a significant spatial signal in the metacommunity and the landscape genetic data (**Fig. 2** and **Table S8** in Supporting information), the asymmetric AEM-based spatial components had a significant pure effect on explaining variation in both community composition as well as genetic composition (**Fig. 2** and **Fig. 3** and **Table S8** in Supporting information). The asymmetric spatial components explained 10% of the variation in the Cladocera metacommunity (F_2,11_=1.907, p=0.04) and 24% of the variation in landscape genetic data (F_2,11_=2.936, p=0.01; **Fig. 2**). The amount of variation in genetic identity explained by environment (total suspended solids) was much lower (7%, F_1,11_= 2.050, p=0.07) than both the amount explained by spatial predictors in the same dataset (24%) and the amount explained in the metacommunity by the same environmental gradient (25%, F_1,11_= 5.265, p=0.001).

**Figure 2.**
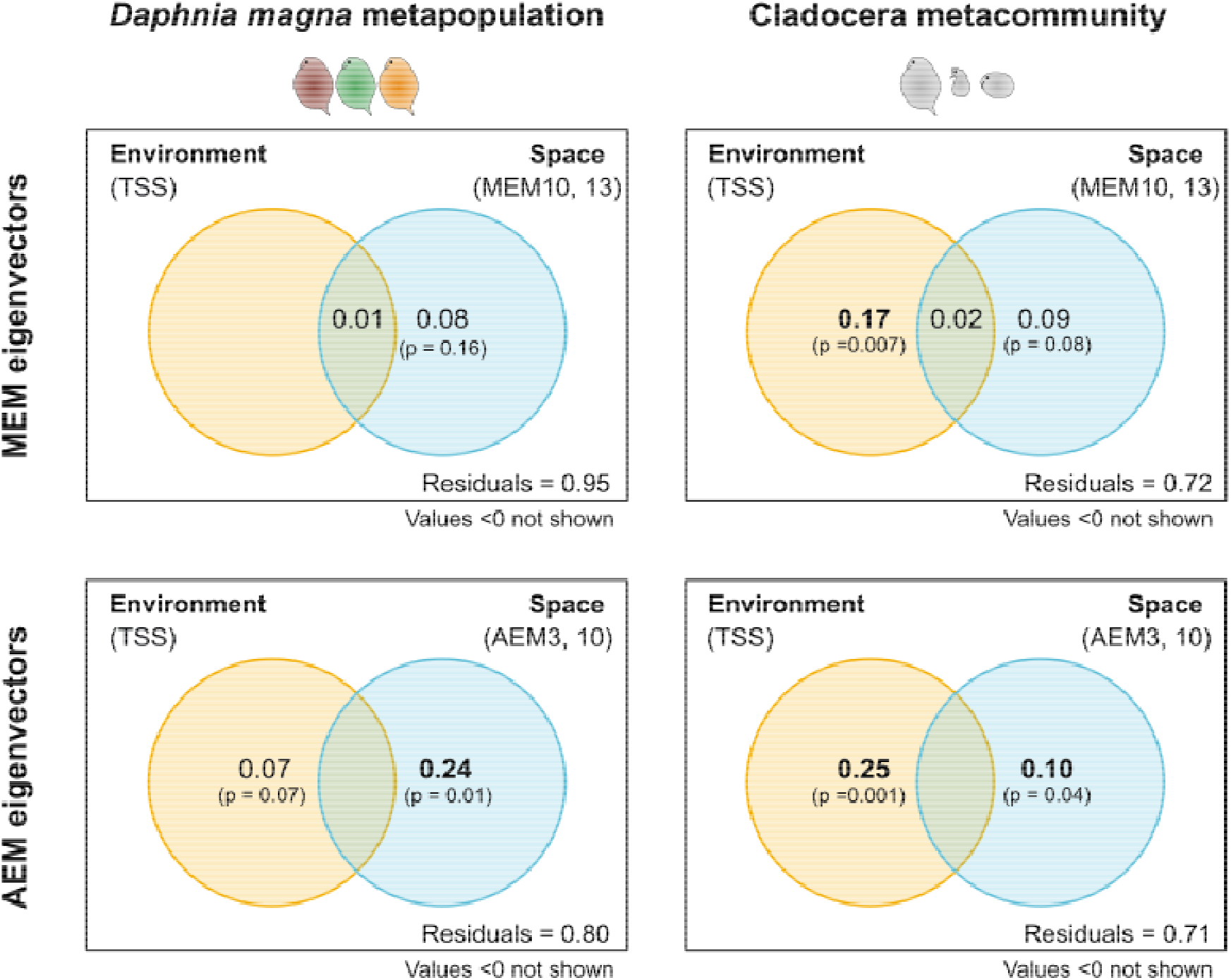
– Variation partitioning results obtained for the Cladocera metacommunity and the *Daphnia magna* metapopulation. Environmental effect only included the concentration of total suspended solids (TSS) in all cases. Spatial predictors were selected from eigenvectors of Moran’s Eigenvector Maps (MEM) for symmetric and Asymmetric Eigenvector Maps (AEM) for wind-related patterns. Bold letters indicate significant pure effects of environment or space (p<0.05). Results of the F-statistics are given in Table S8.

**Figure 3.**
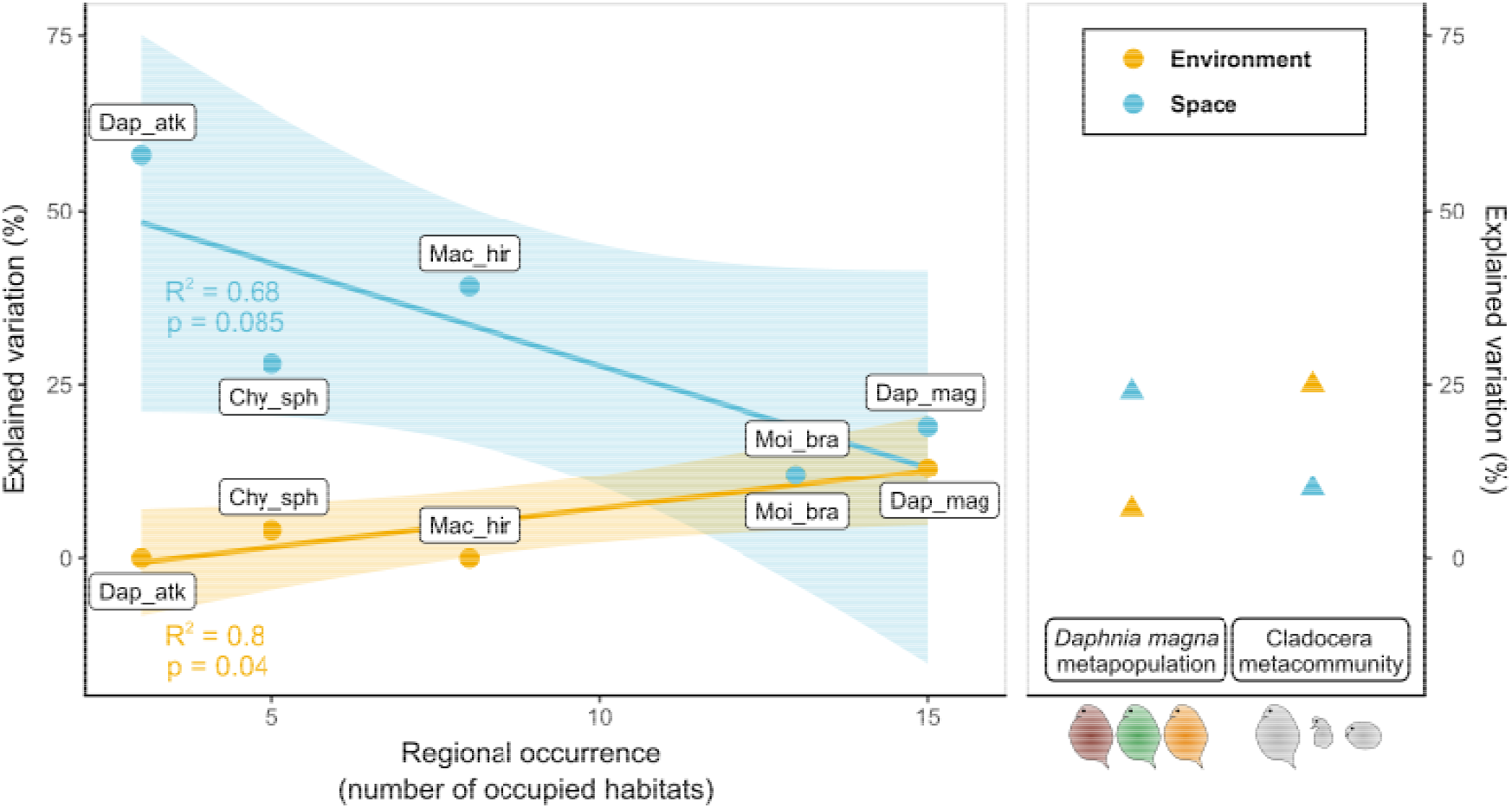
– Left panel: Pure environmental and spatial component of explained variation in the abundance of each member of the Cladocera metacommunity (“Dap_atk” – *Daphnia atkinsoni*, “Chy_sph” – *Chydorus sphaericus*, “Mac_hir” – *Macrothrix hirsuticornis*, “Moi_bra” – *Moina brachiata*, “Dap_mag” – *Daphnia magna*), plotted against their regional occurrence. Spatial predictors were selected from eigenvectors of Asymmetric Eigenvector Maps (AEM). Right panel: Pure environmental and spatial component of explained variation calculated for the metacommunity of Cladocera and for the genetic variation in the *D. magna* metapopulation (presented in detail in Figure 2).

We found a significant relationship between the regional occurrence of species and the pure explanatory power of local environment for cladoceran metacommunity structure (**Fig. 3**; R^2^=0.80, F_1,3_= 11.69, p=0.04), with a higher fraction of variation explained by environmental gradients in more widespread species. An opposite trend was found for the relationship between the pure effect of spatial predictors and regional occurrence (**Fig. 3**; R^2^=0.68, F_1,3_= 6.458, p=0.085). No relationship was found between the total amount of explained variation and regional occurrence (R^2^=0.27, F_1,3_= 1.089, p=0.373). In a parallel analysis on alleles (**Fig. S3** in Supporting information), no effect of regional occurrence was found on the amount of variation explained by space or environment. The amount of variation explained by space (variable AEM components depending on the allele; **Fig. S2** in Supporting information) was relatively high for all alleles, whereas environmental signal was overall very low.

In the metapopulation, we found no clear isolation-by-distance patterns. Genetic distances were not related to spatial distances, neither along nor orthogonal to the dominant wind direction (Mantel’s p = 0.23 for along the wind and p = 0.31 for orthogonal distances; **Fig. 4**).

**Figure 4.**
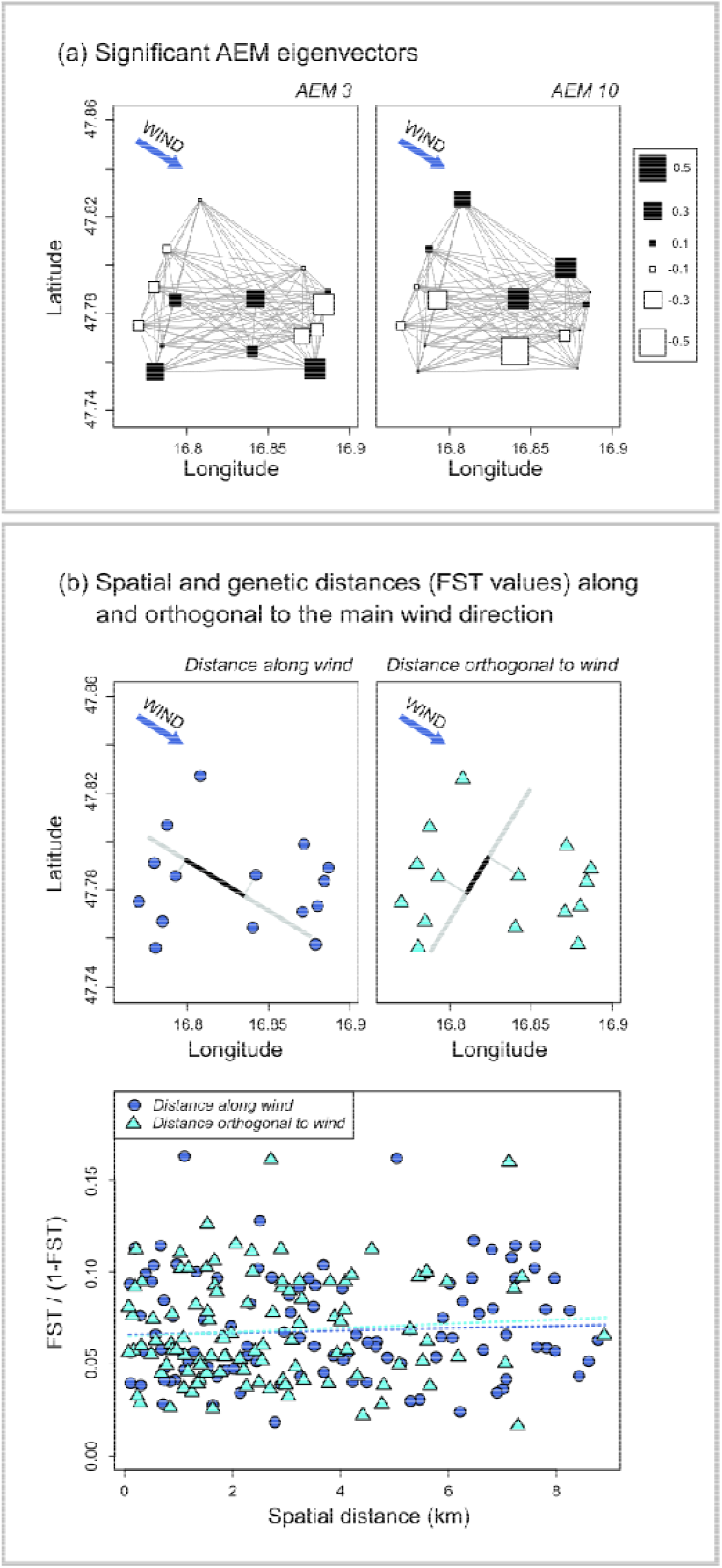
– (a) Eigenvectors of Asymmetric Eigenvector Maps (AEM) retained in the analyses, with their eigenfunction values represented by the size of squares at the spatial position of each habitat. (b) Calculation of spatial distances along and orthogonal to the dominant wind direction (bold lines illustrate a specific example in each case; above) and their relationship with pairwise F_ST_ values (below).

## Discussion

Our comparative analysis highlights clear divergences and some parallels in the relative roles of the main ecological processes in structuring the cladoceran zooplankton metacommunity and the landscape genetic variation of one of its most widespread and abundant species within the same landscape. It shows that while the processes structuring the assembly of communities and populations are similar (Vellend 2010, 2016), they do result in different patterns because of differences in the relative strengths of selection, dispersal or gene flow, and drift in determining the assembly of communities and populations.

Whereas the structure of the metacommunity inhabiting the habitat network is significantly influenced by environmental gradients (total suspended solids), this is not the case for the landscape genetic structure. This may be linked to the fact that microsatellite loci are presumably neutral, and a more detailed genomic analysis might reveal loci and SNPs that do show an association with environmental gradients (e.g., predation, as in Chaturvedi et al. 2021). Yet, our data do indicate that natural selection in this system with a strong salinity gradient is not so strong that it affects the assembly of populations so profoundly that it would affect the structure of neutral genetic variation (as opposed to e.g., hypersaline lakes; Frisch et al. 2021).

A key parallel in metacommunity and landscape genetic structure in the study system is the significant signal of space, and the fact that the explanatory power of space for both metacommunity and metapopulation is linked to the wind directionality captured in our AEM based analysis. For both metacommunity and metapopulation structure, we indeed observed that the amount of variation explained by space is significant in the AEM analysis while not significant in the MEM-based analyses. This indicates that wind is likely an important dispersal agent in this system, in line with the strong winds in the region (Horváth et al. 2016).

The large surface to depth ratio of the soda pans (Boros et al. 2014), their temporary nature exposing sediments and dormant eggs to wind (Vanschoenwinkel et al. 2008; Pinceel et al. 2015; Boros et al. 2017), the strong winds in the region (Horváth et al. 2016), and the presence of large numbers of birds that can act as dispersal agents (Szabó et al. 2022) lead to the expectation that there should be high dispersal rates among habitats in this system. Yet there are some indications of dispersal limitation. First, a historical analysis of species diversity decline driven by the disappearance of rare species with increasing loss of habitat patches suggests non-equilibrium metacommunity structure for these species, reflecting strong abundance-linked dispersal limitation (Horváth et al. 2019). Our data reveal a similar pattern of dispersal limitation, given that the environmental signal increases and the spatial signal declines with increasing abundance of species in the studied metacommunity. Variation in the abundances of *D. magna*, being the most widespread species, is least explained by spatial drivers of all species. This is in contrast, however, with the importance of the spatial signal in explaining the landscape genetic structure of the species. Space is more important in determining landscape genetic structure than in determining occurrences and abundances of this species. This increase in spatial signal is not caused by the presence of many rare alleles that would show a stronger signal of dispersal limitation than more widespread alleles, as there was no association between the spatial signal and the relative frequency of alleles.

While there is a clear spatial signal for the landscape genetic structure, there is no pattern of isolation-by-distance. Isolation-by-distance is a common expectation in population genetics (Slatkin 1987; Bohonak 1999) predicted by theoretical models (Wright 1943; Hartl and Clark 1997) as dispersal (gene flow) is expected to decline with increasing geographic distances. The absence of a pattern of isolation-by-distance has nevertheless also been observed in several other studies (Jenkins et al. 2010; Sexton et al. 2014) including *Daphnia* (Innes 1991; Vanoverbeke and De Meester 1997) and suggests that the processes determining landscape genetic structure deviate from standard expectations reflecting ongoing gene flow, but may for instance also include a signature of patterns of past colonization (Orsini et al. 2013). The spatial signature reflected by the significant contribution of two AEM factors also does not translate into a difference in the relationship between genetic differentiation and geographic distance as measured parallel or orthogonal to the dominant wind direction. This suggests that the impact of wind is differentiated and results in a rather patchy pattern, as it is also reflected in the patterns that are tested for in the significant AEM factors, being AEM3 and AEM10. Wind directionality thus does influence landscape genetic structure (cf. difference between MEM and AEM signals), but in a patchy way. This might reflect that the wind effect is modulated by dispersal mediated by birds. The overall absence of a signal of isolation-by-distance and the patchy nature of the wind effect may reflect that in the soda pans in the Seewinkel region, wind does influence directionality of gene flow, but the strong winds and birds flying from one soda pan to the other might enable equally strong connectivity among pans that are further away from each other than among neighbouring pans. Birds can indeed bridge large distances and do not necessarily move only between nearby patches (Green and Figuerola 2005; Viana et al. 2013; Urgyán et al. 2022). The observed pattern suggests that short- and long-distance dispersal are indeed equally important at the scale of this landscape. At the same time, the level of genetic differentiation (global F_ST_ value of 0.12) is moderate but significant, indicating that gene flow is not so high in this system as to homogenize genetic variation.

Our comparative analysis of metapopulation genetic and metacommunity structure in the same landscape of soda pans reveals some clear divergences (relative role of environmental and spatial drivers) and some parallels (strong indications for a significant contribution of wind directionality to the pattern) between assembly at these two different levels of ecological organisation. We believe that such comparative analyses can contribute to a better understanding of the patterns at each level. In addition, given that population assembly determines evolutionary potential and that patterns of local adaptation may impact community assembly and responses to environmental change (Urban et al. 2008; Hendry 2017; Leibold et al. 2022), the joint analysis of identity and trait variation at the level of populations and communities is important to understand how populations and communities respond to landscape heterogeneity and environmental change (Govaert et al. 2021, 2022).

## Acknowledgements

We thank Adrienn Tóth, Emil Boros, Rudolf Schalli, and Richard Haider for assistance during sample collection, Christian Preiler for laboratory measurements, and Bart Hellemans for practical advice on population genetic analyses. Research was supported by the FWO project G0C3818N, Projekt 06/14216 of the research funding program “International Communication” of ÖFG (Austrian Research Association), the NKFIH project 2019-2.1.11-TÉT-2020-00159, and KU Leuven Research Council project C16/2017/002. ZH acknowledges support from the János Bolyai Research Scholarship of the Hungarian Academy of Sciences and NKFIH FK project 132095.

## References

1. Arar, E. J., and G. B. Collins. 1997. In-vitro determination of chlorophyll a and pheophytin a in marine and freshwater algae by fluorescence: Cincinnati. Ohio, National Exposure Research Laboratory, Office of Research and Development, US Environmental Protection Agency, Method EPA 445.

2. Bivand, R., M. Altman, L. Anselin, R. Assunção, O. Berke, A. Bernat, and G. Blanchet. 2015. Package ‘spdep.’ The Comprehensive R Archive Network.

3. Blanchet, F. G., P. Legendre, and D. Borcard. 2008. Modelling directional spatial processes in ecological data. Ecological Modelling 215: 325–336.

4. Bohonak, A. J. 1999. Dispersal, gene flow, and population structure. The Quarterly Review of Biology 74: 21–45.

5. Bohonak, A. J. 2002. IBD (isolation by distance): a program for analyses of isolation by distance. Journal of Heredity 93: 153–154.

6. Borcard, D., P. Legendre, and P. Drapeau. 1992. Partialling out the Spatial Component of Ecological Variation. Ecology 73: 1045–1055. doi:10.2307/1940179

7. Boros, E., Z. Horváth, G. Wolfram, and L. Vörös. 2014. Salinity and ionic composition of the shallow astatic soda pans in the Carpathian Basin. Annales de Limnologie - International Journal of Limnology 50: 59–69. doi:10.1051/limn/2013068

8. Boros, E., K. V. -Balogh, L. Vörös, and Z. Horváth. 2017. Multiple extreme abiotic conditions and trophic state of intermittent soda pans in the Carpathian Basin (Central Europe). Limnologica 62: 38–46.

9. Chaturvedi, A., J. Zhou, J. A. M. Raeymaekers, and others. 2021. Extensive standing genetic variation from a small number of founders enables rapid adaptation in Daphnia. Nat Commun 12: 4306. doi:10.1038/s41467-021-24581-z

10. Clark, L. V., and M. Jasieniuk. 2011. POLYSAT: an R package for polyploid microsatellite analysis. Mol Ecol Resour 11: 562–566. doi:10.1111/j.1755-0998.2011.02985.x

11. Clesceri, L. S., A. E. Greenberg, and A. D. Eaton. 1999. Standard methods for examination of water & wastewater.

12. Dray, S., G. Blanchet, D. Borcard, and others. 2018. Package ‘adespatial,.’

13. Dray, S., P. Legendre, and P. R. Peres-Neto. 2006. Spatial modelling: a comprehensive framework for principal coordinate analysis of neighbour matrices (PCNM). Ecological Modelling 196: 483–493.

14. Epele, L. B., D. A. Dos Santos, R. Sarremejane, and others. 2021. Blowin’ in the wind: Wind directionality affects wetland invertebrate metacommunities in Patagonia. Global Ecology and Biogeography 30: 1191–1203. doi:10.1111/geb.13294

15. Frantz, A. C., S. Bertouille, M. C. Eloy, A. Licoppe, F. Chaumont, and M. C. Flamand. 2012. Comparative landscape genetic analyses show a Belgian motorway to be a gene flow barrier for red deer (Cervus elaphus), but not wild boars (Sus scrofa). Molecular Ecology 21: 3445–3457. doi:10.1111/j.1365-294X.2012.05623.x

16. Frisch, D., C. Lejeusne, M. Hayashi, M. T. Bidwell, J. Sánchez-Fontenla, and A. J. Green. 2021. Brine chemistry matters: Isolation by environment and by distance explain population genetic structure of Artemia franciscana in saline lakes. Freshwater Biology 66: 1546–1559. doi:10.1111/fwb.13737

17. Goldberg, C. S., and L. P. Waits. 2010. Comparative landscape genetics of two pond-breeding amphibian species in a highly modified agricultural landscape. Molecular ecology 19: 3650–3663.

18. Goudet, J. 2005. Hierfstat, a package for R to compute and test hierarchical F-statistics. Molecular Ecology Notes 5: 184–186.

19. Govaert, L., L. De Meester, S. Rousseaux, S. A. Declerck, and J. H. Pantel. 2021. Measuring the contribution of evolution to community trait structure in freshwater zooplankton. Oikos 130: 1773–1787.

20. Govaert, L., J. H. Pantel, and L. De Meester. 2022. Quantifying eco-evolutionary contributions to trait divergence in spatially structured systems. Ecological Monographs 92: e1531. doi:10.1002/ecm.1531

21. Green, A. J., and J. Figuerola. 2005. Recent advances in the study of long-distance dispersal of aquatic invertebrates via birds. Diversity and Distributions 11: 149–156.

22. Guillot, G., A. Estoup, F. Mortier, and J. F. Cosson. 2005. A spatial statistical model for landscape genetics. Genetics 170: 1261–1280.

23. Hansen, H. P., F. Koroleff, K. Grasshoff, K. Kremling, and M. Ehrhardt. 1999. Methods of seawater analysis. Grasshoff, K., Kremling, K., Ehrhardt, M. Wiley-VCH, Germany.

24. Hartl, D. L., and A. G. Clark. 1997. Principles of population genetics, Sinauer associates Sunderland.

25. Hein, C., H. E. Abdel Moniem, and H. H. Wagner. 2021. Can We Compare Effect Size of Spatial Genetic Structure Between Studies and Species Using Moran Eigenvector Maps? Frontiers in Ecology and Evolution 9.

26. Hendry, A. P. 2017. Eco-evolutionary Dynamics, Princeton University Press.

27. Horváth, Z., R. Ptacnik, C. F. Vad, and J. M. Chase. 2019. Habitat loss over six decades accelerates regional and local biodiversity loss via changing landscape connectance. Ecology Letters 22: 1019–1027.

28. Horváth, Z., C. F. Vad, and R. Ptacnik. 2016. Wind dispersal results in a gradient of dispersal limitation and environmental match among discrete aquatic habitats. Ecography 39: 726–732.

29. Horváth, Z., C. F. Vad, A. Tóth, K. Zsuga, E. Boros, L. Vörös, and R. Ptacnik. 2014. Opposing patterns of zooplankton diversity and functioning along a natural stress gradient: When the going gets tough, the tough get going. Oikos 123: 461–471. doi:10.1111/j.1600-0706.2013.00575.x

30. Horváth, Z., C. F. Vad, L. Vörös, and E. Boros. 2013. The keystone role of anostracans and copepods in European soda pans during the spring migration of waterbirds. Freshwater Biology 58: 430–440. doi:10.1111/fwb.12071

31. Innes, D. J. 1991. Geographic patterns of genetic differentiation among sexual populations of Daphnia pulex. Canadian Journal of Zoology 69: 995–1003.

32. Jenkins, D. G., M. Carey, J. Czerniewska, and others. 2010. A meta-analysis of isolation by distance: relic or reference standard for landscape genetics? Ecography 33: 315–320. doi:10.1111/j.1600-0587.2010.06285.x

33. Jombart, T. 2008. adegenet: a R package for the multivariate analysis of genetic markers. Bioinformatics 24: 1403–1405.

34. Kling, M. M., and D. D. Ackerly. 2021. Global wind patterns shape genetic differentiation, asymmetric gene flow, and genetic diversity in trees. Proceedings of the National Academy of Sciences 118: e2017317118.

35. Leibold, M. A., and J. M. Chase. 2017. Metacommunity ecology, Princeton University Press.

36. Leibold, M. A., L. Govaert, N. Loeuille, L. De Meester, and M. C. Urban. 2022. Evolution and community assembly across spatial scales. Annual Review of Ecology, Evolution, and Systematics 53: 299–326.

37. Leibold, M. A., M. Holyoak, N. Mouquet, and others. 2004. The metacommunity concept: a framework for multi-scale community ecology. Ecology Letters 7: 601–613. doi:10.1111/j.1461-0248.2004.00608.x

38. Manel, S., and R. Holderegger. 2013. Ten years of landscape genetics. Trends in Ecology & Evolution 28: 614–621.

39. Manel, S., M. K. Schwartz, G. Luikart, and P. Taberlet. 2003. Landscape genetics: combining landscape ecology and population genetics. Trends in Ecology & Evolution 18: 189– 197.

40. Manier, M. K., and S. J. Arnold. 2006. Ecological correlates of population genetic structure: a comparative approach using a vertebrate metacommunity. Proceedings of the Royal Society of London B: Biological Sciences 273: 3001–3009.

41. Moncada, B., J. A. Mercado-Díaz, N. Magain, and others. 2021. Phylogenetic diversity of two geographically overlapping lichens: isolation by distance, environment, or fragmentation? Journal of Biogeography 48: 676–689. doi:10.1111/jbi.14033

42. Montero-Pau, J., A. Gómez, and J. Muñoz. 2008. Application of an inexpensive and high-throughput genomic DNA extraction method for the molecular ecology of zooplanktonic diapausing eggs. Limnology and Oceanography: Methods 6: 218–222.

43. Nosil, P., T. H. Vines, and D. J. Funk. 2005. Reproductive isolation caused by natural selection against immigrants from divergent habitats. Evolution 59: 705–719.

44. Oksanen, J., F. G. Blanchet, M. Friendly, and others. 2018. vegan: Community Ecology Package. R package version 2.5–3.

45. Orsini, L., K. I. Spanier, and L. De Meester. 2012. Genomic signature of natural and anthropogenic stress in wild populations of the waterflea Daphnia magna: validation in space, time and experimental evolution. Molecular Ecology 21: 2160–2175.

46. Orsini, L., J. Vanoverbeke, I. Swillen, J. Mergeay, and L. De Meester. 2013. Drivers of population genetic differentiation in the wild: isolation by dispersal limitation, isolation by adaptation and isolation by colonization. Molecular ecology 22: 5983– 5999.

47. Paradis, E. 2010. pegas: an R package for population genetics with an integrated–modular approach. Bioinformatics 26: 419–420.

48. Paradis, E., J. Claude, and K. Strimmer. 2004. APE: analyses of phylogenetics and evolution in R language. Bioinformatics 20: 289–290.

49. Peres-Neto, P. R., P. Legendre, S. Dray, and D. Borcard. 2006. Variation partitioning of species data matrices: estimation and comparison of fractions. Ecology 87: 2614– 2625.

50. Pinceel, T., L. Brendonck, and B. Vanschoenwinkel. 2015. Propagule size and shape may promote local wind dispersal in freshwater zooplankton—a wind tunnel experiment. Limnology and Oceanography.

51. R Development Core Team. 2012. R: A Language and Environment for Statistical Computing.

52. Richardson, J. L. 2012. Divergent landscape effects on population connectivity in two co-occurring amphibian species. Molecular Ecology 21: 4437–4451.

53. Ripley, B., B. Venables, D. M. Bates, K. Hornik, A. Gebhardt, D. Firth, and M. B. Ripley. 2013. Package ‘MASS.’ Cran R 538.

54. Sexton, J. P., S. B. Hangartner, and A. A. Hoffmann. 2014. Genetic isolation by environment or distance: which pattern of gene flow is most common? Evolution 68: 1–15.

55. Slatkin, M. 1987. Gene flow and the geographic structure of natural populations. Science 236: 787–792.

56. Szabó, B., A. Szabó, C. F. Vad, E. Boros, D. Lukić, R. Ptacnik, Z. Márton, and Z. Horváth. 2022. Microbial stowaways: Waterbirds as dispersal vectors of aquatic pro-and microeukaryotic communities. Journal of Biogeography 49: 1286–1298. doi:10.1111/jbi.14381

57. Tóth, A., Z. Horváth, C. F. Vad, K. Zsuga, S. A. Nagy, and E. Boros. 2014. Zooplankton of the European soda pans: fauna, communities and conservation of a unique habitat type. International Review of Hydrobiology 99: 255–276. doi:10.1002/iroh.201301646

58. Truett, G. E., P. Heeger, R. L. Mynatt, A. A. Truett, J. A. Walker, and M. L. Warman. 2000. Preparation of PCR-quality mouse genomic DNA with hot sodium hydroxide and tris (HotSHOT). Biotechniques 29: 52–54.

59. Urban, M. C., M. A. Leibold, P. Amarasekare, and others. 2008. The evolutionary ecology of metacommunities. Trends in Ecology & Evolution 23: 311–317.

60. Urgyán, R., B. A. Lukács, R. Fekete, A. Molnár V., A. Nagy, O. Vincze, A. J. Green, and Á. Lovas-Kiss. 2022. Plants dispersed by a non-frugivorous migrant change throughout the annual cycle. Global Ecology and Biogeography **Early View**. doi:10.1111/geb.13608

61. Vanoverbeke, J., and L. De Meester. 1997. Among-populational genetic differentiation in the cyclical parthenogen Daphnia magna (Crustacea, Anomopoda) and its relation to geographic distance and clonal diversity. Hydrobiologia 360: 135–142. doi:10.1023/A:1003160903708

62. Vanschoenwinkel, B., S. Gielen, M. Seaman, and L. Brendonck. 2008. Any way the wind blows - frequent wind dispersal drives species sorting in ephemeral aquatic communities. Oikos 117: 125–134. doi:10.1111/j.2007.0030-1299.16349.x

63. Vellend, M. 2010. Conceptual synthesis in community ecology. The Quarterly review of biology 85: 183–206.

64. Vellend, M. 2016. The theory of ecological communities, Princeton University Press.

65. Vellend, M., and M. A. Geber. 2005. Connections between species diversity and genetic diversity. Ecology Letters 8: 767–781. doi:10.1111/j.1461-0248.2005.00775.x

66. Viana, D. S., and J. M. Chase. 2019. Spatial scale modulates the inference of metacommunity assembly processes. Ecology 100: e02576. doi:10.1002/ecy.2576

67. Viana, D. S., L. Santamaría, T. C. Michot, and J. Figuerola. 2013. Migratory strategies of waterbirds shape the continental-scale dispersal of aquatic organisms. Ecography 36: 430–438. doi:10.1111/j.1600-0587.2012.07588.x

68. Wang, I. J., and G. S. Bradburd. 2014. Isolation by environment. Mol Ecol 23: 5649–5662. doi:10.1111/mec.12938

69. Weir, B. S., and C. C. Cockerham. 1984. Estimating F-statistics for the analysis of population structure. Evolution 38: 1358–1370.

70. Wright, S. 1943. Isolation by distance. Genetics 28: 114.

