## Supplementary information for "Parallels and divergences in landscape genetic and metacommunity patterns in zooplankton inhabiting soda pans"

**Tables**

**Table S1** – The number of alleles per microsatellite locus in the 15 *Daphnia magna* populations, together with the number of individuals genotyped (Ind), total number of alleles per population (A), mean allelic richness (A_r_), expected (H_e_) and observed heterozygosity across loci (H_o_), and Wright’s inbreeding coefficient (F_IS_). For the habitats, identification code (ID), name, coordinate (latitude and longitude), conductivity value (Cond, in mS/cm) and the concentration of total suspended solids (TSS, in mg/l) are also given. In soda pans, a general multiplying factor of 0.8 can be applied (Boros et al. 2014) between conductivity and salinity (in g/l). Therefore, our conductivity values correspond to 1.4–8.1 g/l salinity.

|  |  |  |  |  |  | **Microsatellite markers** | | | | | | | | | | | | | |  | |  | |  | |  | |  | |
| --- | --- | --- | --- | --- | --- | --- | --- | --- | --- | --- | --- | --- | --- | --- | --- | --- | --- | --- | --- | --- | --- | --- | --- | --- | --- | --- | --- | --- | --- |
| **ID** | **Name of habitat** | **Latitude** | **Longitude** | **Cond** | **TSS** | **A001** | **B008** | **B010** | **B030** | **B045** | **B050** | **B064** | **B074** | **B096** | **B107** | **B133** | **B150** | **B164** | **Ind** | | **A** | | **A_r_** | | **H_o_** | | **H_e_** | | **F_IS_** |
| AS | Albersee | 47.77513639 | 16.77011917 | 8.75 | 837 | 8 | 5 | 4 | 4 | 3 | 11 | 5 | 3 | 4 | 13 | 3 | 6 | 8 | 21 | | 77 | | 5.923 | | 0.50 | | 0.65 | | 0.23 |
| MS | Mittlerer Stinkersee | 47.80675889 | 16.78752194 | 6.19 | 240 | 7 | 7 | 2 | 4 | 4 | 7 | 2 | 5 | 4 | 9 | 2 | 6 | 10 | 20 | | 69 | | 5.308 | | 0.42 | | 0.57 | | 0.26 |
| SM | Sechsmahdlacke | 47.78378861 | 16.88411194 | 4.81 | 1053 | 10 | 6 | 3 | 4 | 4 | 8 | 7 | 4 | 6 | 13 | 2 | 7 | 7 | 20 | | 81 | | 6.231 | | 0.52 | | 0.60 | | 0.24 |
| ÖW | Östliche Wörthenlacke | 47.77334361 | 16.88009389 | 3.25 | 28.8 | 7 | 3 | 3 | 2 | 2 | 7 | 3 | 3 | 5 | 10 | 5 | 9 | 10 | 20 | | 69 | | 5.308 | | 0.47 | | 0.57 | | 0.35 |
| GN | Grosse Neubruchlacke | 47.78612111 | 16.84219833 | 5.52 | 1240 | 9 | 6 | 3 | 2 | 2 | 9 | 7 | 3 | 5 | 12 | 2 | 7 | 8 | 20 | | 75 | | 5.769 | | 0.35 | | 0.59 | | 0.18 |
| NF | Neufeldlacke | 47.76447667 | 16.84024833 | 5.27 | 2320 | 6 | 7 | 4 | 4 | 4 | 8 | 7 | 4 | 4 | 10 | 4 | 6 | 9 | 19 | | 77 | | 5.923 | | 0.42 | | 0.69 | | 0.24 |
| SS | Südlicher Silbersee | 47.79113278 | 16.77973694 | 10.08 | 207 | 6 | 5 | 3 | 6 | 2 | 8 | 7 | 4 | 5 | 11 | 2 | 5 | 10 | 20 | | 74 | | 5.692 | | 0.44 | | 0.58 | | 0.22 |
| ZL | Zicklacke | 47.76696444 | 16.78475194 | 5.1 | 42 | 7 | 7 | 4 | 4 | 2 | 9 | 3 | 4 | 3 | 6 | 5 | 6 | 8 | 20 | | 68 | | 5.231 | | 0.37 | | 0.57 | | 0.13 |
| LL | Lange Lacke | 47.75747528 | 16.87876472 | 3.47 | 800 | 7 | 5 | 2 | 3 | 3 | 6 | 4 | 2 | 5 | 11 | 5 | 9 | 9 | 20 | | 71 | | 5.462 | | 0.50 | | 0.53 | | 0.19 |
| RL | Runde Lacke | 47.78574472 | 16.79274361 | 7.86 | 912 | 11 | 4 | 2 | 3 | 4 | 7 | 6 | 3 | 5 | 10 | 4 | 6 | 8 | 20 | | 73 | | 5.615 | | 0.48 | | 0.59 | | 0.41 |
| KS | Krautingsee | 47.75602861 | 16.78047972 | 4.09 | 540 | 7 | 6 | 5 | 5 | 4 | 10 | 8 | 6 | 6 | 9 | 7 | 8 | 10 | 20 | | 91 | | 7.000 | | 0.55 | | 0.71 | | 0.41 |
| WW | Westliche Wörthenlacke | 47.77092167 | 16.87077667 | 3.88 | 29 | 8 | 5 | 3 | 2 | 5 | 8 | 9 | 3 | 5 | 11 | 6 | 7 | 8 | 20 | | 80 | | 6.154 | | 0.49 | | 0.64 | | 0.06 |
| OH | Obere Höllacke | 47.82711944 | 16.80793222 | 8.68 | 426 | 8 | 4 | 2 | 4 | 3 | 8 | 5 | 3 | 4 | 7 | 2 | 5 | 7 | 10 | | 62 | | 4.769 | | 0.45 | | 0.58 | | 0.24 |
| AL | Auerlacke | 47.78917083 | 16.88677778 | 1.97 | 860 | 11 | 5 | 2 | 4 | 3 | 9 | 4 | 3 | 4 | 7 | 5 | 5 | 8 | 16 | | 70 | | 5.385 | | 0.44 | | 0.64 | | 0.32 |
| SL | Stundlacke | 47.79890389 | 16.87170639 | 1.76 | 314 | 8 | 6 | 3 | 2 | 2 | 8 | 8 | 3 | 4 | 9 | 5 | 4 | 8 | 20 | | 70 | | 5.385 | | 0.45 | | 0.62 | | 0.27 |
|  |  |  |  |  |  | 17 | 10 | 7 | 9 | 8 | 17 | 12 | 9 | 10 | 22 | 10 | 12 | 15 |  | |  | |  | |  | |  | |  |

**Table S2** – Pairwise Pearson's correlations among the environmental predictors (Z – water depth, TSS – concentration of total suspended solids, Chl – concentration of chlorophyll-a, TP – concentration of total phosphorus) in the 15 habitats. Upper triangle: p values, lower triangle: r values.

|  | **Z** | **TSS** | **conductivity** | **Chl** | **TP** |
| --- | --- | --- | --- | --- | --- |
| **Z** |  | 0.025 | 0.828 | 0.131 | 0.048 |
| **TSS** | -0.576 |  | 0.593 | 0.001 | 0.000 |
| **conductivity** | -0.062 | 0.150 |  | 0.807 | 0.458 |
| **Chl** | -0.408 | 0.764 | -0.069 |  | 0.001 |
| **TP** | -0.517 | 0.802 | 0.208 | 0.757 |  |

**Table S3** – Significant environmental predictors explaining the variation in each Cladocera species in the region, according to AIC-based permutations

| **Species** | **Z** | **TSS** | **conductivity** | **Chl** |
| --- | --- | --- | --- | --- |
| *Daphnia atkinsoni* | ● |  | ● |  |
| *Chydorus sphaericus* |  | ● |  |  |
| *Macrothrix hirsuticornis* | ● |  |  |  |
| *Moina brachiata* |  |  | ● | ● |
| *Daphnia magna* | ● | ● | ● | ● |

**Table S4** – Significant spatial predictors (AEM eigenvectors; see also **Fig. S3**) explaining the variation in each Cladocera species in the region, according to AIC-based permutations

| **Species** | **1** | **2** | **3** | **4** | **5** | **6** | **7** | **8** | **9** | **10** | **11** |
| --- | --- | --- | --- | --- | --- | --- | --- | --- | --- | --- | --- |
| *Daphnia atkinsoni* | ● | ● | ● | ● | ● | ● | ● | ● |  | ● | ● |
| *Chydorus sphaericus* |  |  |  |  | ● | ● |  |  | ● |  | ● |
| *Macrothrix hirsuticornis* | ● |  |  | ● | ● | ● |  | ● | ● | ● | ● |
| *Moina brachiata* | ● | ● | ● | ● | ● | ● | ● | ● | ● |  | ● |
| *Daphnia magna* | ● |  |  |  |  |  |  |  |  | ● | ● |

**Table S5** – Results of the variation partitioning for each Cladocera species in the region (used in **Figure 3** in the main text)

| **Species** | **Regional occurrence** | **Pure env (%)** | **Shared (%)** | **Pure space (%)** | **Unexplained (%)** |
| --- | --- | --- | --- | --- | --- |
| *Daphnia atkinsoni* | 3 | 0 | 40 | 58 | 2 |
| *Chydorus sphaericus* | 5 | 4 | 26 | 28 | 42 |
| *Macrothrix hirsuticornis* | 8 | 0 | 29 | 39 | 34 |
| *Moina brachiata* | 13 | 12 | 40 | 12 | 34 |
| *Daphnia magna* | 15 | 13 | 34 | 19 | 37 |

**Table S6** – Observed (H_o_) and expected heterozygosity (H_e_) and F-statistics (F_IS_, F_IT_, and F_ST_) for all loci across 15 soda pan populations. F_IS_: mean deficit of heterozygotes within populations; F_IT_: mean global deficit of heterozygotes across populations; F_ST_: mean fixation index of each population.

| **Microsatellite** | **H_o_** | **H_e_** | **F_IS_** | **F_IT_** | **F_ST_** |
| --- | --- | --- | --- | --- | --- |
| **A001** | 0.54 | 0.74 | 0.23948994 | 0.2794589 | 0.05255547 |
| **B008** | 0.52 | 0.64 | 0.14925629 | 0.1921104 | 0.05037251 |
| **B010** | 0.36 | 0.44 | 0.01868972 | 0.1920727 | 0.17668522 |
| **B030** | 0.14 | 0.56 | 0.65708493 | 0.7472341 | 0.26289073 |
| **B045** | 0.22 | 0.55 | 0.59660609 | 0.6052739 | 0.02148710 |
| **B050** | 0.72 | 0.84 | 0.11269366 | 0.1482893 | 0.04011653 |
| **B064** | 0.63 | 0.79 | 0.10568377 | 0.2013783 | 0.10700303 |
| **B074** | 0.18 | 0.59 | 0.61504723 | 0.6957169 | 0.20955730 |
| **B096** | 0.39 | 0.72 | 0.34607012 | 0.4659726 | 0.18335675 |
| **B107** | 0.62 | 0.88 | 0.27837455 | 0.3038462 | 0.03529757 |
| **B133** | 0.3 | 0.41 | 0.20456550 | 0.2783614 | 0.09277435 |
| **B150** | 0.66 | 0.74 | 0.01887353 | 0.1124320 | 0.09535820 |
| **B164** | 0.68 | 0.87 | 0.18382106 | 0.2206815 | 0.04516220 |

**Table S7** – Pairwise F_ST_ values (lower triangle) and significance levels (upper triangle) among all soda pan populations. The code used for each pan is given in **Table S1** (as ID).

|  | **AS** | **MS** | **SM** | **ÖW** | **GN** | **NF** | **SS** | **ZL** | **LL** | **RL** | **KS** | **WW** | **OH** | **AL** | **SL** |
| --- | --- | --- | --- | --- | --- | --- | --- | --- | --- | --- | --- | --- | --- | --- | --- |
| **AS** | - | 0.005 | 0.002 | 0.000 | 0.007 | 0.000 | 0.003 | 0.000 | 0.000 | 0.000 | 0.003 | 0.000 | 0.003 | 0.000 | 0.000 |
| **MS** | 0.036 | - | 0.064 | 0.000 | 0.002 | 0.000 | 0.000 | 0.000 | 0.000 | 0.000 | 0.000 | 0.000 | 0.002 | 0.000 | 0.000 |
| **SM** | 0.044 | 0.026 | - | 0.000 | 0.005 | 0.000 | 0.000 | 0.000 | 0.000 | 0.000 | 0.000 | 0.000 | 0.000 | 0.000 | 0.000 |
| **ÖW** | 0.058 | 0.093 | 0.096 | - | 0.000 | 0.000 | 0.000 | 0.004 | 0.000 | 0.000 | 0.000 | 0.000 | 0.000 | 0.000 | 0.000 |
| **GN** | 0.032 | 0.044 | 0.040 | 0.054 | - | 0.000 | 0.002 | 0.000 | 0.000 | 0.000 | 0.000 | 0.000 | 0.002 | 0.000 | 0.000 |
| **NF** | 0.061 | 0.095 | 0.096 | 0.075 | 0.080 | - | 0.000 | 0.000 | 0.000 | 0.000 | 0.000 | 0.000 | 0.000 | 0.000 | 0.000 |
| **SS** | 0.047 | 0.097 | 0.105 | 0.057 | 0.064 | 0.077 | - | 0.003 | 0.000 | 0.022 | 0.002 | 0.000 | 0.000 | 0.000 | 0.000 |
| **ZL** | 0.048 | 0.093 | 0.105 | 0.037 | 0.058 | 0.054 | 0.039 | - | 0.000 | 0.000 | 0.000 | 0.000 | 0.000 | 0.000 | 0.000 |
| **LL** | 0.090 | 0.141 | 0.142 | 0.089 | 0.090 | 0.115 | 0.063 | 0.090 | - | 0.021 | 0.000 | 0.000 | 0.000 | 0.000 | 0.000 |
| **RL** | 0.047 | 0.104 | 0.107 | 0.063 | 0.059 | 0.091 | 0.030 | 0.055 | 0.031 | - | 0.003 | 0.000 | 0.000 | 0.000 | 0.000 |
| **KS** | 0.040 | 0.068 | 0.075 | 0.058 | 0.066 | 0.060 | 0.042 | 0.064 | 0.064 | 0.044 | - | 0.006 | 0.000 | 0.000 | 0.000 |
| **WW** | 0.042 | 0.053 | 0.056 | 0.057 | 0.049 | 0.079 | 0.072 | 0.080 | 0.056 | 0.050 | 0.036 | - | 0.269 | 0.042 | 0.003 |
| **OH** | 0.041 | 0.048 | 0.055 | 0.087 | 0.051 | 0.092 | 0.082 | 0.090 | 0.065 | 0.059 | 0.052 | 0.020 | - | 0.005 | 0.003 |
| **AL** | 0.061 | 0.074 | 0.073 | 0.073 | 0.063 | 0.086 | 0.095 | 0.090 | 0.071 | 0.076 | 0.051 | 0.029 | 0.041 | - | 0.004 |
| **SL** | 0.056 | 0.053 | 0.056 | 0.088 | 0.051 | 0.086 | 0.103 | 0.100 | 0.105 | 0.088 | 0.076 | 0.041 | 0.046 | 0.042 | - |

**Table S8** – Details including the F-statistics of the variation partitioning analyses reported in **Figure 2** and **3** in the main text

| Response dataset | Spatial predictors | Explanatory dataset | Adjusted R^2^ | F-value | Between-groups degrees of freedom | Within-groups degrees of freedom | p-value |
| --- | --- | --- | --- | --- | --- | --- | --- |
| *Daphnia magna* metapopulation | MEM | Environment | -0.05 | 0.414 | 1 | 11 | 0.83 |
|  |  | Space | 0.08 | 1.566 | 2 | 11 | 0.16 |
|  | AEM | Environment | 0.07 | 2.050 | 1 | 11 | 0.07 |
|  |  | Space | 0.24 | 2.936 | 2 | 11 | 0.01 |
| Cladocera metacommunity | MEM | Environment | 0.17 | 3.872 | 1 | 11 | 0.007 |
|  |  | Space | 0.09 | 1.811 | 2 | 11 | 0.08 |
|  | AEM | Environment | 0.25 | 5.265 | 1 | 11 | 0.001 |
|  |  | Space | 0.10 | 1.907 | 2 | 11 | 0.04 |

**Figures**

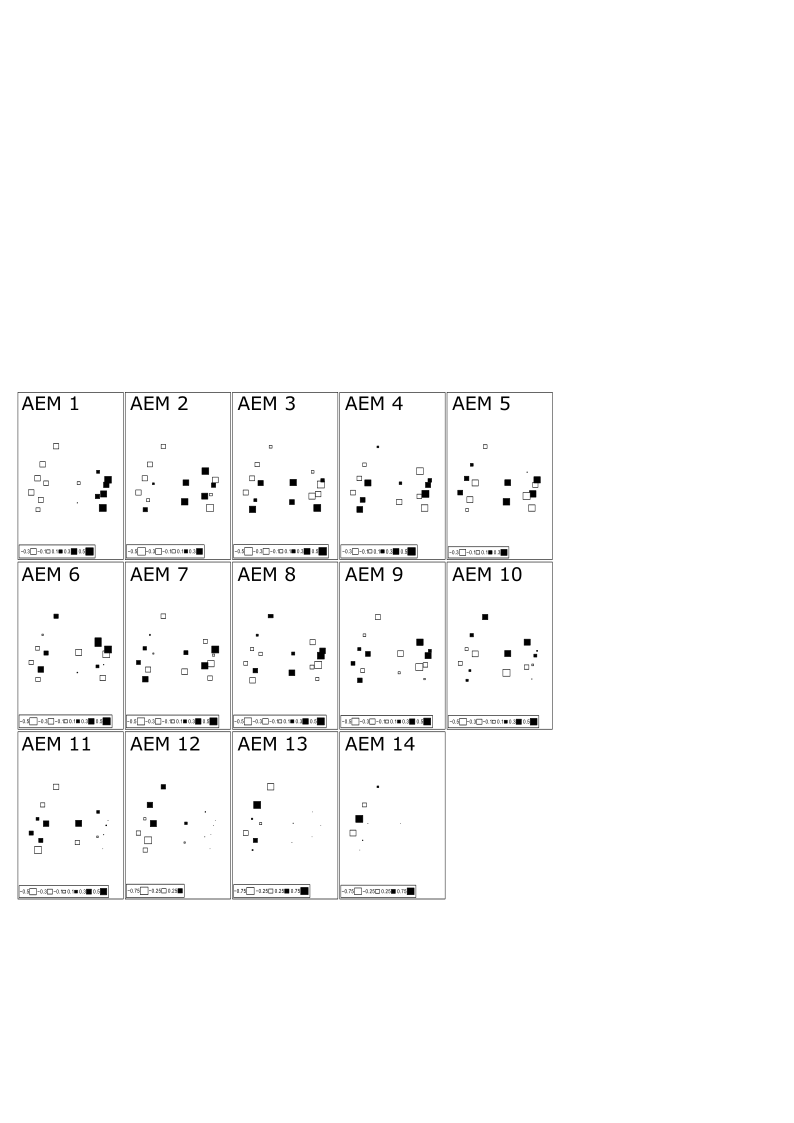
**Figure S1**– All eigenvectors of Asymmetric Eigenvector Maps (AEM) used to capture the spatial distances and directional links of the 15 habitats in the network, with their eigenfunction values represented by the size of squares at the spatial position of each habitat. AEM 3 and AEM 10 were retained in the model selections, explaining a general north-south (AEM 3) and a northeast-southwest split (AEM 10) in spatial similarities

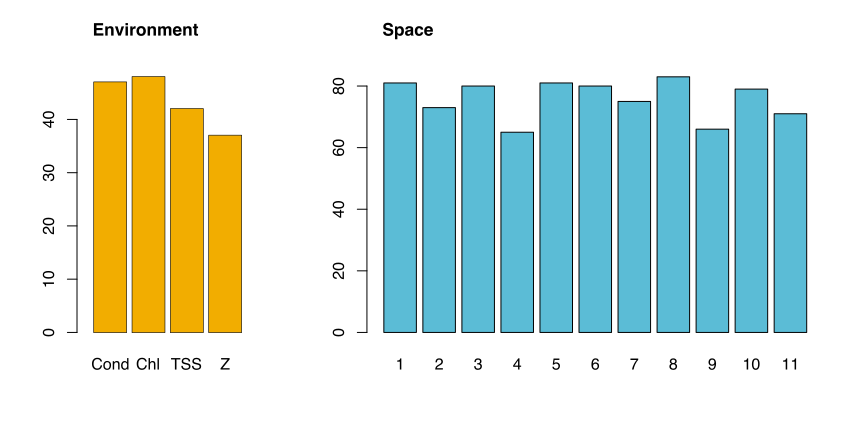

**Figure S2** – The number of times each environmental parameter (left) and AEM eigenvector (right) was retained in AIC-based model selections and used in the variation partitioning analyses ran separately for the 131 alleles that had at least two regional occurrences. Abbreviations: Cond – conductivity, Chl – chlorophyll *a* concentration, TSS – concentration of total suspended solids, Z – water depth; AEM eigenvectors are numbered.

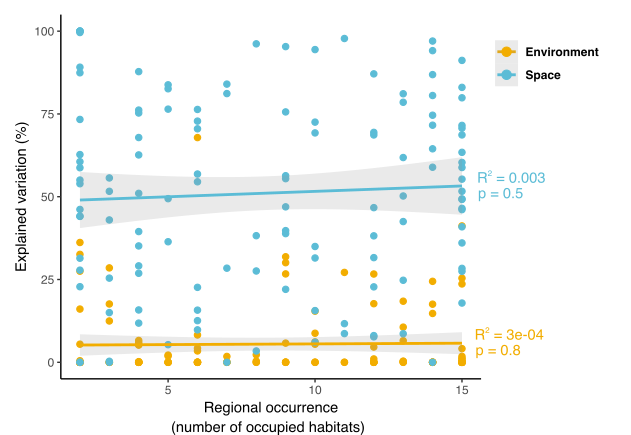

**Figure S3 –** Pure environmental and spatial component of explained variation in the abundance of each allele in the *Daphnia magna* metapopulation, plotted against their regional occurrence. (F-statistics of the linear regressions: F_1,124_=0.042 for the environmental, F_1,124_=0.369 for the spatial model)
